# Development of a modified weight-drop apparatus for closed-skull, repetitive mild traumatic brain injuries in a mouse model

**DOI:** 10.1101/2025.07.25.666403

**Authors:** Anthony B. Crum, Cara D. Nielson, Kaylin J. Sevilla Lopez, Julian P. Meeks

## Abstract

Repetitive mild traumatic brain injury (rmTBI) is a major contributor to long-term neurological dysfunction, yet many preclinical models lack precise control and quantification of biomechanical forces across impacts. We developed a reproducible, closed-skull mouse model of rmTBI using a custom-built weight-drop apparatus featuring a solenoid-based rebound arrest system, integrated high-speed videography, and accelerometry to track head kinematics during impact. Adult male and female mice received either a single impact or nine daily impacts. Linear and angular acceleration data were analyzed alongside behavioral and histological outcomes. Our apparatus delivered consistent impact and velocity forces with minimal inter-subject variability. Additionally, the animals experienced consistent linear and angular acceleration as measured using high-speed video capture. These impacts did not cause skull fracture or acute vascular hemorrhage, but impacted animals had increased return of righting reflex (RoRR) time, consistent with mild, concussion-like symptoms. Behavioral testing revealed reduced performance of rmTBI-affected mice in an olfaction-mediated foraging task (buried food task), particularly at later timepoints, consistent with progressive olfactory impairment. Immunohistochemical analysis of Iba1 and CD68 in the brain demonstrated sex-dependent microglial activation, with males showing higher expression levels in both single- and nine-impact models. Among the brain regions investigated, microglial activation was most pronounced in the corpus callosum, neocortex, and olfactory tubercle. These findings underscore the importance of including sex as a biological variable in rmTBI research and support the utility of this model for probing injury thresholds, regional vulnerability, and potential therapeutic interventions in repetitive head trauma.

**Significance Statement:** Mild traumatic brain injuries (mTBIs) contribute long-term sensory, motor, and cognitive dysfunction. We developed a novel approach for delivering repetitive mTBIs (rmTBIs) to mice via a custom weight-drop apparatus. The device allows precise control over impact forces and enables quantification of linear and angular acceleration during each impact. We describe the apparatus, the forces delivered, and the kinematics experienced by lightly anesthetized mice. We measured behavioral and neuroinflammatory sequelae in the brains of rmTBI-exposed mice compared to controls. rmTBI-exposed animals showed impairment in the olfaction-mediated buried food task and evidence of microglial reactivity in multiple brain regions days-to-weeks following injury. The results demonstrate the utility of this approach for studying rmTBI-associated pathophysiology, and for testing therapies or interventions for rmTBI.

## Introduction

Mild traumatic brain injury (mTBI), often referred to as a concussion, is one of the most common yet underrecognized neurological injuries worldwide. Despite being classified as “mild,” these injuries can result in persistent physical, emotional, cognitive, and sensory disturbances. Estimates suggest over 2.5 million TBI-related emergency department visits occur annually in the United States, with as many as 75 – 90% of these considered mild in severity (Cassidy et al., 2004; Miller et al., 2021). However, these figures likely underestimate the true incidence, as many mTBIs go unreported due to misdiagnosis, limited public awareness, or lack of consistent clinical guidelines (Bazarian et al., 2005; Langlois et al., 2006).

Repetitive mild TBIs (rmTBI), which may occur in athletic, military, or civilian populations exposed to recurrent low-level impacts, pose an even greater concern. Unlike moderate or severe injuries, rmTBI often eludes detection by conventional imaging, yet has been linked to long-term deficits in cognition, emotional regulation, and sensory processing, including anosmia (Doty et al., 1997; Fann et al., 2004; Proskynitopoulos et al., 2016; Mouzon et al., 2018; Lecuyer Giguere et al., 2019). Accumulating evidence suggests that even subconcussive impacts, particularly when repetitive, can have lasting effect on brain function and structure (Bazarian et al., 2014; Mouzon et al., 2014; Brody et al., 2015; Hirad et al., 2019; Kulkarni et al., 2019)

Mechanistically, mTBI arises from a complex interplay between linear and angular acceleration forces that induce diffuse biomechanical stress within the brain (Meaney and Smith, 2011). These forces can cause microstructural damage, particularly diffuse axonal injury (DAI), along with blood-brain barrier disruption, neuroinflammation, and long-term neurodegenerative changes (Mac Donald et al., 2007; Johnson et al., 2013). Angular acceleration, in particular, has been identified as a critical biomechanical component in the pathogenesis of TBI, contributing disproportionately to white matter injury (Meaney et al., 1995; Owens et al., 2018). Yet, many preclinical models have room for improvement in adequately capturing these kinematics.

This translational gap underscores the need for refined animal models capable of replicating human-relevant TBI biomechanics with precision and reproducibility. Existing rodent models, such as fluid percussion (FPI) and controlled cortical impact (CCI), often lack control over angular acceleration or fail to preserve closed-skull conditions (Dixon et al., 1991; Osier and Dixon, 2016; Carlson et al., 2017). Weight-drop models offer a noninvasive, scalable alternative but are frequently hindered by inconsistent impact delivery, variable rebound, and limited kinematic monitoring (Kane et al., 2012). The blast injury model replicates the overpressure wave dynamics associated with explosive blasts and has helped illuminate TBI mechanisms relevant to military settings; however, it requires complex setups and does not generalize well to non-blast civilian injuries (Long et al., 2009). The laboratory of Dr. Cheryl Wellington developed the Closed-Head Impact Model of Engineered Rotational Acceleration (CHIMERA) model, which causes both linear and angular acceleration under controlled, closed-head conditions (Namjoshi et al., 2017; Vonder Haar et al., 2019). CHIMERA uses a pneumatic actuator to generate mTBIs that cause linear and angular acceleration and is compatible with both single and multiple-injury paradigms (Namjoshi et al., 2014). The forces generated by single CHIMERA impacts induce neuroinflammation, minutes-long loss of righting reflex, and behavioral deficits, which are labeled “mild” TBIs but might be consistent with a moderate-to-strong concussion (Namjoshi et al., 2017). Our custom-modified weight-drop apparatus was designed with the goal of delivering mTBIs with linear and angular components, but which allow the delivery of repeated (>2) concussions milder than those produced by CHIMERA. We accomplished this goal by integrating real-time accelerometry, high-speed videography, and an active solenoid-based rebound arrest system. This setup enables reproducible, closed-skull injury delivery while capturing the biomechanical signatures critical to understanding rmTBI pathophysiology. We evaluated the behavioral impacts of these injuries using the buried food task, an olfaction-mediated natural foraging task (Alberts and Galef Jr, 1971; Doty et al., 1997). Given the high prevalence of olfactory deficits in human mTBI and their potential to serve as early biomarkers of injury or recovery (De Luca et al., 2023), this behavioral endpoint offers a translationally relevant readout of functional impairment. To probe cellular responses to injury, we further employed quantitative immunohistochemistry for Iba1 and CD68, markers of microglial activation and phagocytic activity. Microglial reactivity has been implicated in both the acute and chronic phases of mTBI pathogenesis, with sex-specific differences increasingly recognized as important modifiers of injury outcomes (Witcher et al., 2015; Scott et al., 2022).

Through this integrative approach, linking biomechanical, behavioral, and cellular outcomes, our model provides a robust and scalable platform to study the complex pathophysiology of rmTBI. Importantly, this system enables future exploration of injury thresholds, temporal dynamics of inflammation, and therapeutic interventions with high translational relevance.

## Methods

### Construction of modified weight-drop apparatus

A custom weight-drop device was constructed using 80/20 T-slot aluminum framing (80/20 Inc., Columbia City, IN), inspired by the design used by the Peter Douglas Laboratory at the University of Texas Southwestern (Solano Fonseca et al., 2021). The frame includes a central vertical rail for sled guidance, flanked by two lateral rails and three horizontal support bars. Structural stability was reinforced by four 45° angled braces connecting the vertical supports to a rectangular base, which was outfitted with six adjustable leveling feet (Fig. 1A). A lab jack (C & A Scientific, Sterling, VA) mounted on the central support bar was used to position the foam impact platform at the desired height relative to the mouse’s skull.

**Figure 1.**
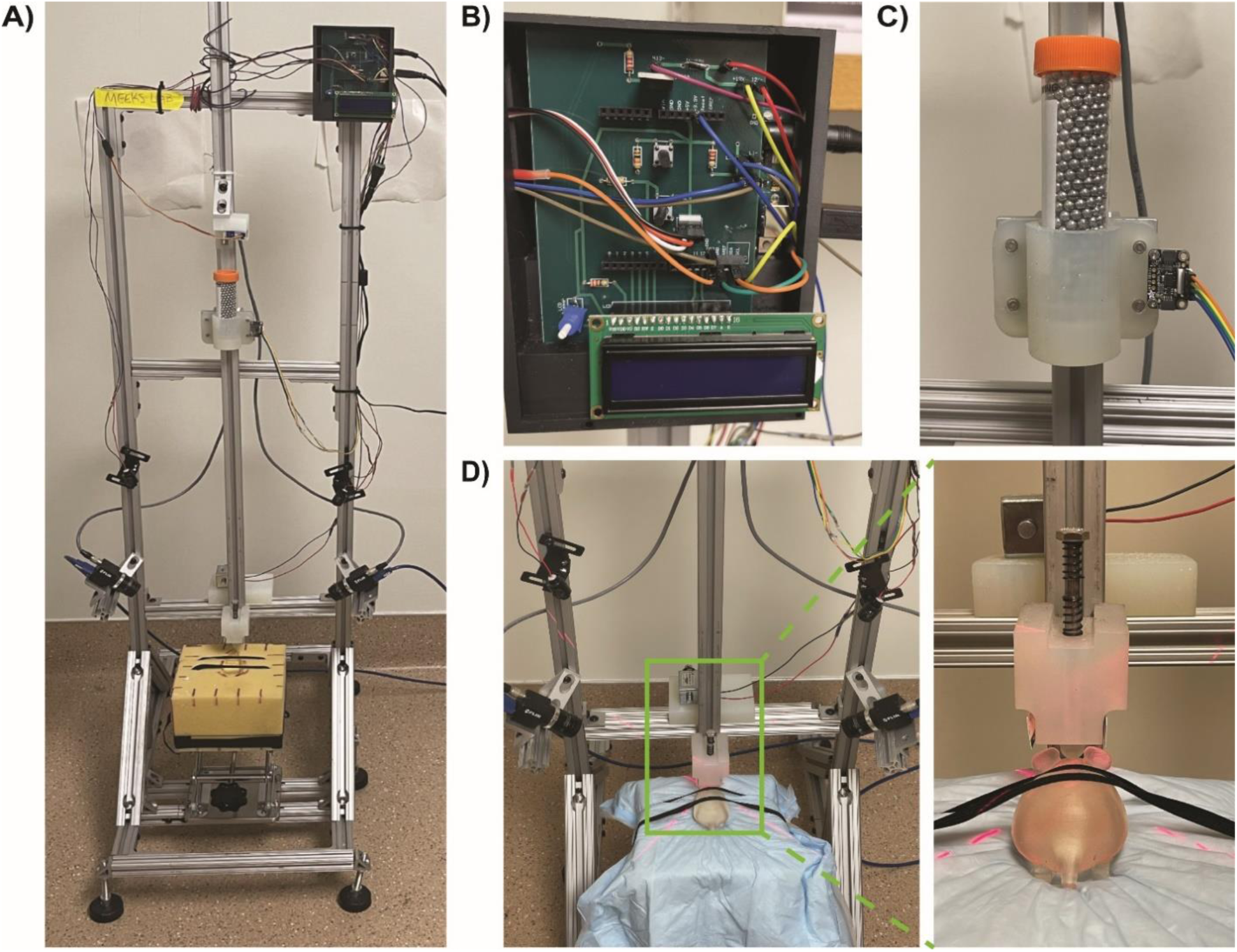
Construction of a Modified Weight-Drop Device: A) Fully assembled modified weight-drop device. B) Close-up view of the custom control module, which includes a custom printed circuit board (PCB), an Arduino Uno v3, and an LCD screen. C) Close-up view of the weight sled, composed of a conical tube filled with steel airgun shot, a custom-printed conical holder, and an accelerometer. D) Close-up view of the high-speed camera placement alongside the solenoid, impactor bolt, and positioning lasers, presented with a 3D-printed mouse model on a foam pad. The inset shows a close-up view of the mouse’s position with the impactor tip above bregma.

Custom components—including mounts for the impactor, servo, solenoid, and weight sled— were designed and 3D-printed using high-durability resin (Formlabs Inc., Somerville, MA). A MicroServo SG90 9g servo motor (Tower Pro) within the servo mount controlled the release of the weight sled via a locking pin mechanism. The sled consisted of a 50 mL conical tube (CELLTREAT Scientific Products, Ayer, MA; Product #229421) filled with 4.5 mm steel airgun pellets (Daisy Outdoor Products, Rogers, AR), housed in a 3D-printed holder, and attached to an 80/20 15 Series 3-slot carriage (Part #6535) that slides along the vertical rail (Fig. 1C).

To ensure precise head positioning beneath the impactor, two red laser diodes (Adafruit Industries, Brooklyn, NY; Product #1054) were mounted on opposing sides of the frame to project an intersecting crosshair above the bregma suture (Fig. 1D). The impactor consisted of a ¼-inch diameter, ½-inch long steel bolt fitted with a spring to induce rebound and capped with a nylon tip to evenly distribute force.

Two Blackfly S USB3 cameras (Teledyne FLIR LLC, Wilsonville, OR) were mounted laterally to capture high-speed video for subsequent kinematic analysis. An Arduino Uno Rev3 microcontroller (Arduino, Somerville, MA), integrated with a custom printed circuit board (ExpressPCB LLC, Mulino, OR), managed servo actuation, accelerometer data acquisition, and solenoid triggering (Fig 1B). This modular design enhanced stability, functionality, and ease of component replacement.

### Induction of repetitive mild traumatic brain injury

This device attached an accelerometer to a weighted impact “sled” in order to log the acceleration of the sled before, during, and after it contacted a steel impactor with a blunted nylon tip. We determined the maximum combination of weights and heights for the sled empirically which, in recently euthanized mice, did not cause skull fracture or acute vascular hemorrhage. These conditions were further validated in a pilot cohort of deeply (ketamine/xylazine) anesthetized mice, establishing the parameters utilized in this study.

Eight-week-old male and female C57BL/6J mice (The Jackson Laboratory, Bar Harbor, ME) in the single-impact cohort received a single subcutaneous injection of 1.0 mg/kg extended-release buprenorphine (Wedgewood Pharmacy, Swedesboro, NJ) for pre-procedural analgesia. Animals in the nine-impact rmTBI cohort received a subcutaneous injection of 1.0 mg/kg extended-release buprenorphine prior to each set of three impacts for a total of three doses. Animals in both cohorts were then anesthetized using a 3% isoflurane/97% oxygen mixture in a VetFlo™ induction chamber (Kent Scientific Corporation, Torrington, CT).

Each mouse was positioned on a sterile pad placed over a foam block and secured with two elastic straps (Fig. 1D). The impactor tip was aligned above the bregma suture, in contact with the intact scalp. Upon initiation, the Arduino code activated the servo, rotating its arm from 90° to 170° to remove the locking pin, allowing the 444.35 g weight sled to drop from a height of 0.5 meters. The impactor holder restricts the displacement of each impact to 2 cm, which pressed the head into the foam base. For multiple impact studies, following the impact, a spring surrounding the impactor bolt induced sled rebound, and when the onboard ADXL345 accelerometer recorded a change in the directionality of the acceleration, the software triggered the solenoid (Adafruit Industries; Product #413) to engage and arrest the sled, preventing secondary impacts.

### Quantification of impact force

The ADXL345 accelerometer recorded acceleration approximately every 15 milliseconds. Acceleration values were plotted over time, and the value immediately preceding the directional reversal was taken as the peak impact acceleration. Instantaneous velocity at impact was calculated using the kinematic equation, v = u + aΔt, where v is final velocity, u is initial velocity, a is acceleration, and Δt is the elapsed time. Impact force was then calculated using the impulse-momentum relationship, F = mv / d, where m is the sled mass (0.44435 kg), v is velocity at impact, and d is the impact displacement (0.02 m).

### Quantification of linear and angular acceleration

High-speed videos from each impact event were analyzed using Kinovea motion tracking software (Charmant, 2024). Digital markers were placed on anatomical and device landmarks, including the nose tip, right ear midpoint, bottom of the impactor, bottom of the frame, and bottom of a fixed ruler. Frame-by-frame tracking ensured accurate capture of linear and angular displacement across time. Videos without clear visual representation of any one of the anatomical landmarks were excluded from analysis. The resulting kinematic data were exported and analyzed in JMP® Pro (Version 17.0.0, SAS Institute Inc., Cary, NC).

### Buried Cookie Task

To assess olfactory function, a modified buried food task was conducted, based on protocols from Alberts and Galef (Alberts and Galef Jr, 1971). There were two different cohorts of mice, a cohort for the single-impact model and a cohort for the nine-impact repetitive model. Mice in the single impact cohort were evaluated at baseline (∼8 weeks of age), and again at one, three, and seven days after injury. Prior to testing, animals were familiarized with honey-flavored Teddy Grahams (Mondelēz International, Chicago, IL) to eliminate novelty bias. Mice in the nine-impact repetitive cohort were evaluated at baseline (∼8 weeks of age), and again at one day after the third impact, one day after the sixth impact, one day after the ninth impact, one week after the ninth impact, and one month after the ninth impact. Prior to testing in the nine-impact cohort, mice were familiarized with Annie’s Organic Honey Bunny Grahams (General Mills, Inc., Minneapolis, MN). On test days, mice were habituated for 10 minutes in a clean cage with fresh bedding. The animal was removed from the cage, and the cookie buried ∼2 cm beneath the bedding in a cage corner. Animals were reintroduced into the center of the cage and video-recorded using a Panasonic HC-V100 camera (Panasonic North America, Newark, NJ). Latency to uncover the cookie was recorded, with a 600-second maximum trial duration. Mice were returned to their home cages after each trial and were allowed to consume the cookie regardless of task success.

### Immunohistochemistry

Eight days post-final impact, mice were euthanized via cardiac perfusion with 4% paraformaldehyde (PFA). Brains were post-fixed in 4% PFA for 72 hours, followed by sequential immersion in 20% and 30% sucrose solutions until fully equilibrated. Each brain was bisected sagittally, embedded in Fisher Healthcare Tissue-Plus O.C.T. Compound (Thermo Fisher Scientific, Waltham, MA), and frozen on dry ice. Serial sagittal sections (25 μm) were obtained using a Shandon FSE cryostat (Thermo Fisher Scientific, Waltham, MA).

Tissue sections were selected based on anatomical integrity and inclusion of target regions implicated in TBI pathology (e.g., hippocampus, olfactory bulb, cerebellum). Immunostaining was performed using anti-CD68 (1:1000, Bio-Rad Laboratories, Hercules, CA, Cat. No. MCA1957T) and anti-Iba1 (1:1000, FUJIFILM Wako Pure Chemical Corp., Osaka, Japan, Cat. No. 019-19741) primary antibodies. Secondary antibodies included Alexa Fluor 488-conjugated goat anti-rat IgG (1:1000, Invitrogen, Waltham, MA, Cat. No. A-11006) and Alexa Fluor 633-conjugated goat anti-rabbit IgG (1:1000, Invitrogen, Waltham, MA, Cat. No. A-21070). To visualize cell nuclei, slices were stained with DAPI (1:1000, Thermo Fisher Scientific, Waltham, MA Cat. No. D1306). Fluorescent imaging was conducted at 20X magnification using a Keyence BZ-X800 epifluorescence microscope (Keyence Corporation, Itasca, IL).

For the single-impact TBI model, images were processed in ilastik (Berg et al., 2019), a supervised machine learning platform, to segment and quantify CD68^+^ cells, resting and activated microglia, and double-positive cells. Regions of interest (ROIs) were defined for the accessory olfactory bulb (AOB), cerebellum (CB), cortex (CTX), hindbrain (HB), hippocampal formation (HPF), main olfactory bulb (MOB), olfactory tubercle (OT), piriform cortex (PIR), and substantia nigra (SN). Normalized cell counts were analyzed using least squares regression models in JMP Pro.

For the nine-impact rmTBI model, images were again processed in ilastik to segment and quantify CD68^+^ puncta, low-intensity Iba1^+^ signal (representing broader microglial field presence), high-intensity Iba1^+^ signal (reflecting dense microglial cell bodies), and Iba1^+^ processes (capturing microglial arborization). The resulting binary masks were used to assess the effects of sex and treatment on IHC marker expression across brain regions using a generalized linear mixed model (GLMM) with a binomial distribution and a logit link function. The dependent variable was the proportion of pixels above the 0.5 probability threshold (i.e., predicted as positive). This approach accounts for the non-normal distribution and bounded nature (0–1) of proportion data. Sample ID was included as a random effect to account for repeated measures across regions and to capture inter-subject variability. GLMMs were chosen to account for the hierarchical structure of the data and to avoid violating assumptions of normality and homoscedasticity. ROIs were defined for the nucleus accumbens (ACB), AOB, anterior olfactory nucleus (AON), CB, corpus callosum (CC), caudoputamen (CP), CTX, HB, HPF, lateral olfactory tract (LOT), MOB, optic tract (OPT), OT, PIR, and SN.

## Results

### Establishing conditions for mTBI production with the modified weight-drop device

We recreated the basic architecture of a weight-drop device developed in previous studies (Flierl et al., 2009; Khalin et al., 2016; Chakraborty et al., 2021), and established a set of conditions (weights, heights of sled) for the device that produce the maximal impact forces that do not produce skull fracture or acute vascular hemorrhage (see Materials and Methods). A major goal of our study was to ensure that mTBIs produced were reliable and capable of being accurately quantified. Previous implementations of this weight-drop device produced only simple estimates of impact forces and did not allow detailed analysis of impact dynamics, which are critical when assessing trauma (Kimpara and Iwamoto, 2012; Bodnar et al., 2019). We therefore modified the device to enable active logging of accelerometer data, which we used to track the acceleration and velocities of the sled during each impact time course (Fig. 2A-B).

**Figure 2.**
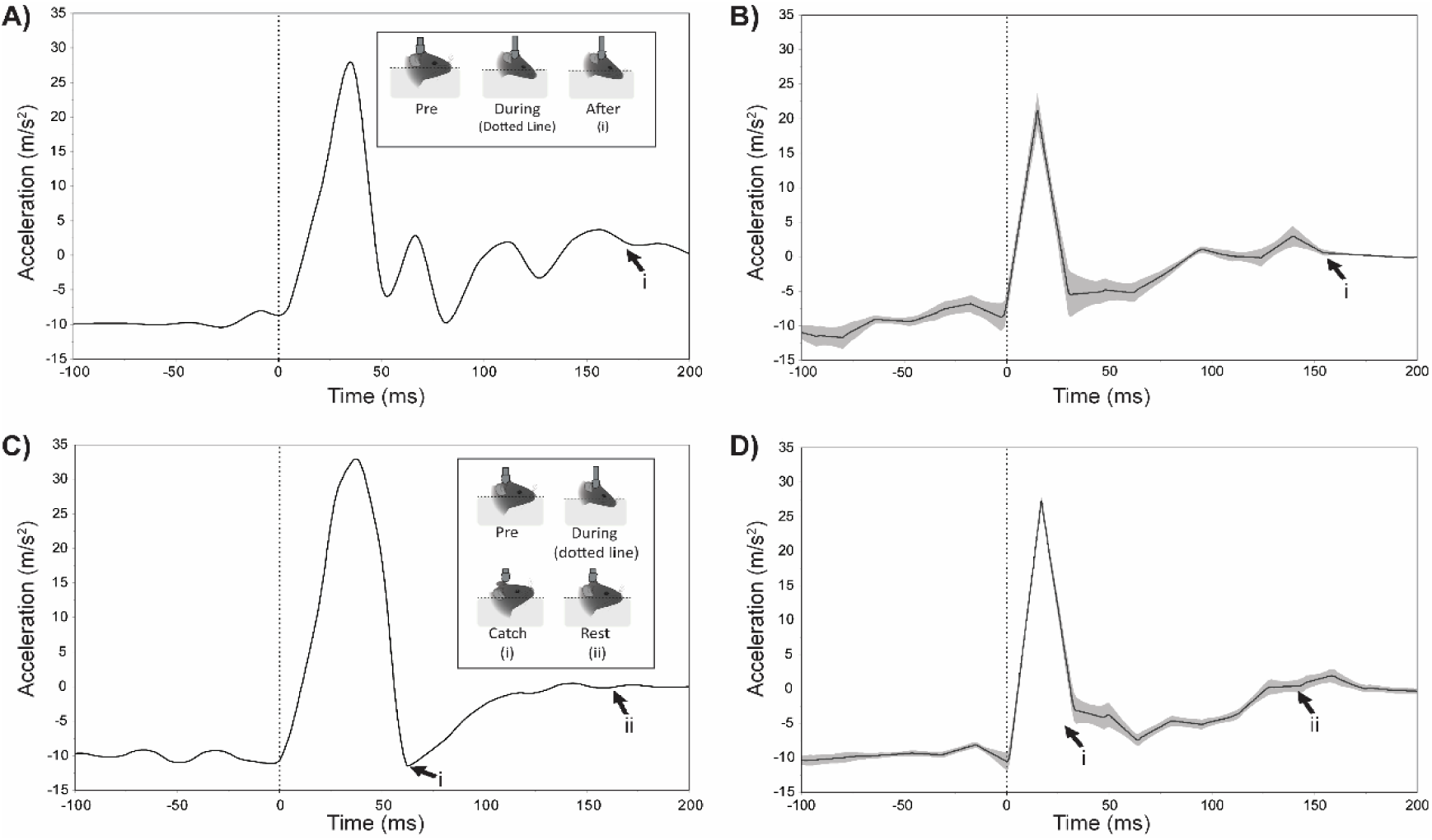
Impactor acceleration profiles following head impact in single- and repetitive-injury models. **(A, B)** Impactor acceleration in the 1-hit model, shown as a representative trial (A) and group average ± SE (B), aligned with the moment of impact (dotted line at 0 ms). **(C,D)** Impactor acceleration in the 9-hit model, which used a solenoid-controlled restraint to prevent secondary impacts, shown as individual trial **(C)** and group average ± SE **(D)**. Inserts illustrate key postural phases: Pre, During, and After impact. In **(A, B)**, **(i)** marks the moment when the weight comes to rest on the animal’s head following impact. In **(C, D)**, **(i)** marks the moment the weight sled contacts the solenoid, and **(ii)** marks when the weight comes to rest on the solenoid arm, effectively preventing secondary contact with the head. Inserts for **(A)** and **(C)** were created using BioRender. Confidence intervals for **(B)** and **(D)** are indicated by the shaded area.

In a cohort of 10 adult male and 10 female C57Bl6\J mice, we quantified critical parameters of each impact. We measured the peak impact velocity, which was 2.44 ± 0.03 m/s (n = 20). We found no difference in the peak impact velocity between male and female mice (2.43 ± 0.05 m/s and 2.44 ± 0.05 m/s, respectively; one-way ANOVA, *F*(1,18) = 0.022, *p* = 0.884). A pooled *t*-test further validated the lack of sex differences in peak impact velocity (*t*(18) = 0.15, *p* = 0.884, 95% CI: [−0.136, 0.156]). Because the weight and height of the sled was the same for each impact, we expected the device to produce consistent impact forces, which we confirmed. Again, no significant difference was found between sexes (one-way ANOVA *F*(1,18) = 0.019, *p* = 0.892), with mean forces of 66.19 ± 2.63 N for females and 66.70 ± 2.63 N for males. The pooled *t*-test supported this finding, *t*(18) = −0.14, *p* = 0.892, 95% CI: [−8.33, 7.30]. Additionally, we measured the recovery of righting reflex (RoRR)—a common measure for return of consciousness (Hamm, 2001)—which was 14.6 ± 3.56 s (n = 40). A two-way ANOVA showed no significant main effect of sex (*F*(1,36) = 2.296, *p* = 0.139), but did indicate a trend-level effect of treatment (*F*(1, 36) = 3.694, *p* = 0.063) with control mice averaging 7.75 ± 5.04 s and injured mice averaging 21.45 ± 5.04 s. The model reported no significant interaction between sex and treatment (*F*(1, 36) = 2.787, *p* = 0.104). These results confirm that the device produces consistent impacts on adult mice of both sexes.

### Enhancing weight-drop device to induce whiplash and monitor linear and angular acceleration

In the initial implementation of the weight-drop device the impactor drove the mouse skull into the foam cushion substrate, where it remained until the experimenter retracted the impactor and removed the mouse from the device (Solano Fonseca et al., 2021). This implementation succeeded in causing a compression injury, but failed to produce significant whiplash effects, which are common in mTBIs and a critical contributor to the stress placed on brain components (Meaney and Smith, 2011). We therefore further modified the weight drop device to allow the mouse’s head to rebound from its maximally compressed position, creating a whiplash effect (Fig. 2C-D). We accomplished this by installing a stiff spring around the impactor bolt, and adding a computer-controlled solenoid piston that restrained the weight sled after impact, preventing a second contact with the impactor (Fig. 1). Following the modifications, we again evaluated the mean peak impact velocity and impact forces which were 3.06 ± 0.02 m/s (95% CI: [3.03, 3.09]) and 104.4 ± 1.01 N (95% CI: [102.4, 106.4]) respectively (n = 99 impacts, 6 male and 5 female mice), both a slight increase over the original implementation. We attribute these changes to subtle modifications made during the upgrade, for example freshly printed parts and improved rail lubrication. Again, we saw no difference in the peak impact velocity between male and female mice (3.05 ± 0.02 m/s and 3.07 ± 0.02 m/s respectively; one-way ANOVA *F*(1, 97) = 0.282, *p* = 0.597) or peak impact force between male and female mice (103.91 ± 1.37 N and 104.97 ± 1.51 N respectively; one-way ANOVA *F*(1, 97) = 0.272, *p* = 0.603). As before, the impacts did not cause skull fractures or vascular hemorrhage. Additionally, we analyzed the RoRR for the 9-impact cohort and found an average of 54.83 ± 3.85 s (95% CI: [47.19, 62.48]) for all animals. We then performed a 3-way ANOVA to determine if there were statistically significant main effects of sex, treatment, impact number, or their interactions. The model indicated a statistically significant effect of treatment, *F*(1, 67) = 4.439, *p* = 0.039 with control animals exhibiting an average RoRR of 46.51 ± 5.59 s (95% CI: [35.35, 57.67]) and injured animals exhibiting an average RoRR of 64.65 ± 5.24 s (95% CI: [52.19, 73.10]). This represents a significant difference between the single impact and 9-impact models and suggests a cumulative effect on return to consciousness in the rmTBI model. Furthermore, this result verifies the change in RoRR is due to the injury status and not an artifact caused by general anesthesia. All other effects were statistically non-significant with *p*-values > 0.2.

In order to quantify the whiplash effects introduced by our modifications, we installed high-speed video cameras to record mouse head motion during each impact (Fig. 3). We tracked points on the mouse head using kinematic software and measured the absolute peak linear and angular accelerations which were 4.28 ± 0.13 m/s^2^ (95% CI: [4.02, 4.55]) and 9,692.84 ± 445.78 rad/s^2^ (95% CI: [8,807.08, 10,578.61]), respectively (88 impacts, 6 male and 5 female mice). We observed no difference in peak linear acceleration between male and female mice (4.28 ± 0.18 m/s^2^ and 4.29 ± 0.20 m/s^2^ respectively; one-way ANOVA *F*(1, 86) = 0.002, *p* = 0.964), or in peak angular acceleration between male and female mice (10,299.25 ± 593.60 rad/s^2^ and 8,934.84 ± 663.67 rad/s^2^ respectively; one-way ANOVA *F*(1, 86) = 2.348, *p* = 0.129).

**Figure 3.**
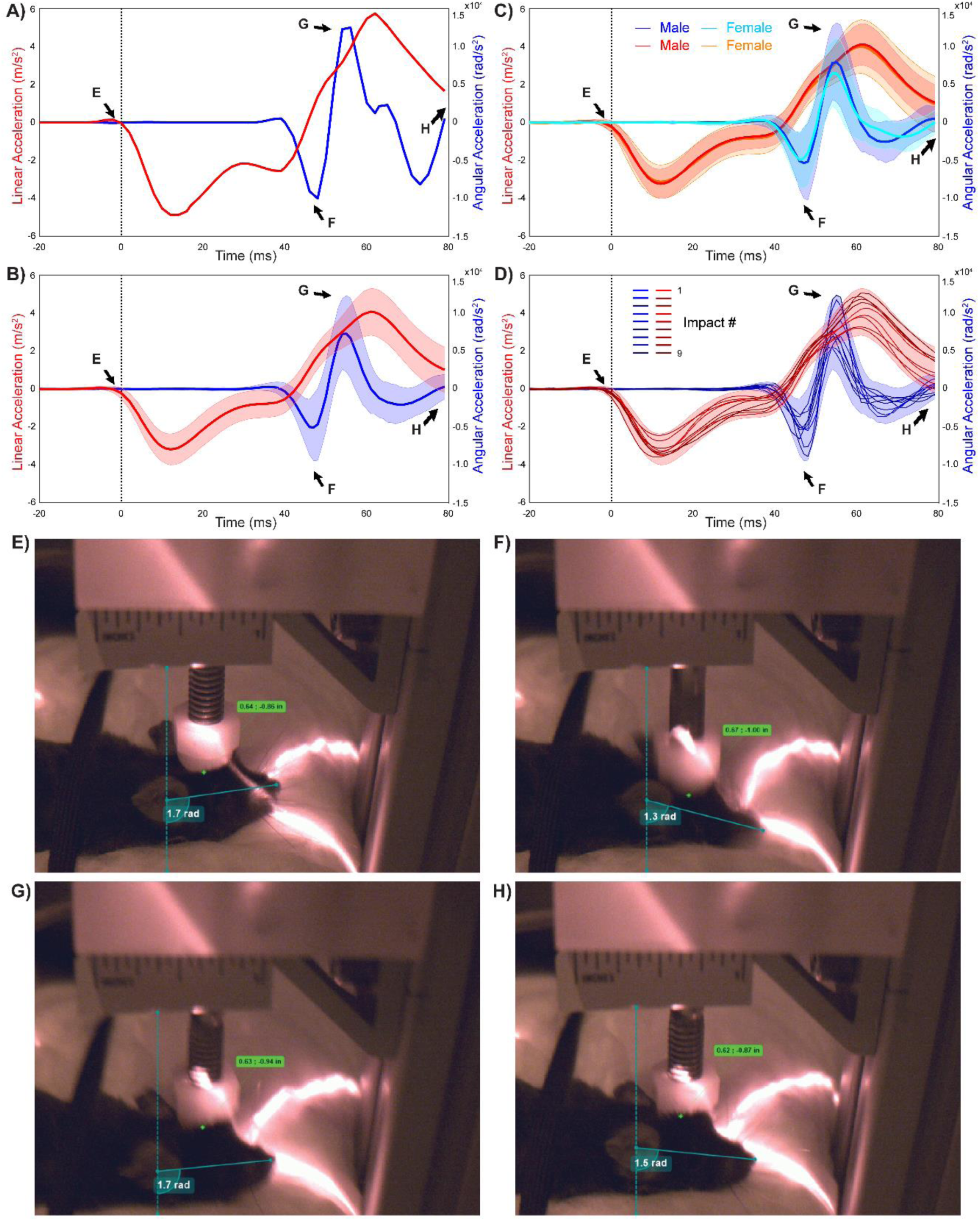
Kinematic profiles for the 9-hit rmTBI model with annotated video frame references. **(A)** Representative linear (red) and angular (blue) acceleration traces from a single trial, aligned to impact (0 ms, vertical dotted line). Arrows labeled E-H correspond to times of video frames shown in Panes E-H. **(B)** Group average across all impacts (mean ± standard deviation). **(C)** Group averages (mean ± standard deviation) for each sex. **(D)** Group averages per impact. Darker shades reflect impacts later in the sequence. The shaded area reflects the standard deviation across all impacts (from Panel B). (**E-H**) Still frames with annotations made in Kinovea, showing angular displacement (radians) and linear distance. The specific frames shown reflect: **(E)** pre-impact rest, **(F)** peak downward angular acceleration, **(G)** peak upward angular acceleration (rebound), and **(H)** return to rest post-oscillation. Tracking points include the bottom frame border, ruler base, right ear midpoint, nose tip, and bottom of the impactor cap.

### Establishing a nine-hit rmTBI protocol including whiplash

The improvements to the device achieved our goals of producing reliable, quantifiable impacts in male and female mice that recapitulate the main components of mTBI. In observing the animals after the injuries in the home cage and general olfactory behavior tasks (to be described later in the manuscript), we noted that all animals tolerated single mTBIs with minimal deleterious effects over the course of days-to-one-week. This observation, combined with knowledge that humans experiencing rmTBIs have higher rates of neurological and neuropsychiatric conditions (Mouzon et al., 2014; Montenigro et al., 2017; Mouzon et al., 2018; Kulkarni et al., 2019), led us to evaluate the use of this device in a nine-impact rmTBI protocol. For this rmTBI protocol, we used the updated version of the weight-drop device including the whiplash component and high-speed video monitoring. We delivered 9 impacts in batches, with 3 impacts per week spaced no less than 24 hours apart, across 3 weeks (see Materials and Methods). This approach was selected because it allowed animals a period of recovery between each impact and gave a 3 – 4 day interval between impact batches to evaluate animal behavior (reported later in the manuscript).

We assessed the performance of the device and the reliability of the resulting injuries in the nine-impact protocol as discussed above, including impact velocity, impact force, linear acceleration, and angular acceleration. For this assessment, we were particularly interested in evaluating the potential for batch effects (*i.e.*, differences in the performance on different experimental days) and sex-specific differences. Peak impact velocity measured via accelerometer logging showed no significant differences across impact sessions (linear mixture model, *F*(8, 72 = 1.645, *p* = 0.127) or the sex of the animal (linear mixture model, *F*(1, 9) = 0.338, *p* = 0.576). Similarly, we found no significant interaction between impact number and sex (linear mixture model, *F*(8, 72) = 1.137, *p* = 0.349). We also found no significant effect of Subject ID (linear mixture model, estimate = −0.00027, SE = 0.0011, 95% CI = [−0.0023, 0.0018], Wald *p* = 0.8027), indicating that none of the animals received a systematically higher or lower impact velocity with the instrument.

The same statistical comparison applied to impact forces delivered by the instrument similarly revealed no effects. We found no main effect of impact number (linear mixture model *F*(8, 72) = 1.624, *p* = 0.133), sex (linear mixture model *F*(1, 9) = 0.335, *p* = 0.577), or an interaction between impact number and sex (linear mixture model *F*(8,72) = 1.135, *p* = 0.351). The contribution of Subject ID was again negligible (Estimate = −1.55, SE = 4.71, 95% CI = [−10.78, 7.68], Wald *p* = 0.742), further confirming the lack of systematic differences across individuals. Collectively, these results confirm the reliability of the device in a rmTBI protocol involving nearly 100 specific impacts spaced across 9 experimental sessions and 3 weeks’ time.

Having confirmed that the instrument itself performed reliably in these conditions, we proceeded to evaluate the “animal side” of the equation – whether male and female animals experienced reliable impacts as assessed by kinematic analysis of linear and angular acceleration. For this analysis, we evaluated both the peak downward linear and angular acceleration (during the compression phase, negative values) and peak upward linear and angular acceleration (during the rebound phase, positive values, see Fig. 3). In contrast to what was observed on the “instrument side,” on the “animal side” we found some noteworthy differences in these measurements.

For maximum “downward” linear acceleration (during the compression phase), we observed no main effect of sex (linear mixture model *F*(1, 10.2) = 0.005, *p* = 0.947), but did observe a significant main effect of impact number (linear mixture model *F*(8, 63.3) = 3.137, *p* = 0.005). Post-hoc tests indicated a significantly increased downward linear acceleration on Impact #7 (Difference = −1.131 ± 0.266 m/s^2^, 95% CI:[−1.889, −0.374]], *p* < 0.0001) and significantly reduced downward linear acceleration on Impact #6 (Difference = 0.544 ± 0.266 m/s^2^, 95% CI: [−0.213, 1.302], *p* = 0.045). The statistical interaction between sex and impact number was not significant (linear mixture model *F*(8, 63.3) = 1.726, *p* = 0.110). The variance component associated with Subject ID (individual animals’ experience of maximum downward acceleration across impacts) accounted for 3.62% of the total model variance (Estimate = 0.030, SE = 0.062, 95% CI: [−0.091, 0.151], Wald *p* = 0.627), indicating minimal inter-subject variability in downward linear acceleration.

For maximum “upward” linear acceleration (during the rebound/whiplash phase), we observed no main effect of sex (linear mixture model, *F*(1, 9.4) = 0.019, *p* = 0.894), but did observe a main effect of impact number (linear mixture model, *F*(8, 62.3) = 4.483, *p* = 0.0002) with the mean of Impact #4 being slightly below the overall average and the mean of Impact #7 being slightly above the overall average. Furthermore, the variance associated with Subject ID accounted for 12.2% of the total model variance (Estimate = 0.149, SE = 0.136, 95% CI: [−0.117, 0.415], Wald *p* = 0.273), indicating modest inter-subject effects. Overall, upward linear acceleration terms showed signs of modest effects across impact number (the “batch” of impacts delivered on specific days), without a pronounced sex bias, which may be partially accounted for by inter-subject variability.

Finally, we performed complimentary analysis on maximum downward (in the clockwise direction with the mouse head facing to the right) and upward (in the counterclockwise direction) angular acceleration across sexes and impact numbers. For maximum downward angular acceleration, we observed no statistically significant effects of sex (linear mixture model *F*(1, 9.4) = 2.075, *p* = 0.182), impact number (linear mixture model *F*(8, 70) = 1.550, *p* = 0.156), or interaction between sex and impact number (linear mixture model *F*(8, 70) = 1.051, *p* = 0.408), and the variance associated with Subject ID was negligible (Estimate =98,165.765, SE = 666,060, 95% CI: [−1,207,288, 1,403,619.4], Wald *p* = 0.883).

For maximum upward angular acceleration (angular acceleration during rebound/whiplash phase), we observed no significant effect of impact number (linear mixture model *F*(8, 70) = 1.446, *p* = 0.193). We did observe a significant main effect of sex across impacts (linear mixture model *F*(1, 10.4) = 5.403, *p* = 0.042). Specifically, female mice experienced decreased upward angular acceleration during whiplash (Difference = −1,745.92 ± 751.105 rad/s^2^, 95% CI: [−3,411.05, −80.800], *p* = 0.042)). There was no statistically significant interaction between sex and impact number (linear mixture model *F*(8, 70) = 1.555, *p* = 0.155). The variance associated with Subject ID was again negligible (Estimate =-814,711.2, SE = 655,304.9, 95% CI: [−2,099,085, 469,662.8], Wald *p* = 0.214). Collectively, these results revealed that this paradigm can introduce mild batch effects (associated with experimental days), which may be attributable to minor differences in the positioning of the foam block, placement of the animals, or other details of the impact procedure. These batch effects resulted in no systematically weaker or stronger forces, or head kinematic changes to any individual animal in the cohorts. Interestingly, maximal upward angular acceleration was lower in females than in males across impacts, suggesting they experienced modestly lower whiplash forces in this paradigm.

### Evaluating behavioral changes in an olfaction-associated foraging task (Buried Cookie Task)

In both the single-impact (no whiplash) and nine-impact (with whiplash) variants of the mTBI paradigm, we evaluated the effects of these injuries on mouse behavior using a simple olfaction-associated foraging task (the Buried Cookie Task, see Materials and Methods). This task was chosen for two main reasons: (1) it is simple to implement and quantify and (2) the task is heavily – though not entirely – dependent on intact olfactory function, a particular interest of the laboratory. A third minor reason was that the model, when applied across successive sessions, may allow for analysis of motor, attention, and learning components. These components might be evaluated in the future with the assistance of recent advancements in machine learning-assisted behavioral analysis (Mathis et al., 2018; Wang et al., 2023; Correia et al., 2024). Here, we focused our analysis on the most common measurements made from buried food task trials: Retrieval Time and Retrieval Success.

In the single-impact paradigm, retrieval time (the time before an animal finds the buried cookie, maximum 600 seconds) and retrieval success (a binary success/failure value) were measured at four timepoints: a pre-treatment baseline and post-treatment timepoints at 1, 3, and 7 days following the injury (Fig. 4). Control animals were anesthetized, placed on the mTBI apparatus, but received no impact prior to recovering from anesthesia. We observed no significant effect of time point (linear mixture model, *F*(3, 108) = 1.981, *p* = 0.121) or treatment (linear mixture model, *F*(1, 36) = 0.536, *p* = 0.469). However, we did observe a significant fixed effect of sex (*F*(1, 36) = 6.798, *p* = 0.0132), with females exhibiting longer retrieval times than males (Estimate = +74.888, SE = 28.722, 95% CI: [16.637, 133.138], *p* = 0.013). We also observed a significant interaction between sex and timepoint (linear mixture model *F*(3, 108) = 2.732, *p* = 0.047), with males demonstrating reduced retrieval times (finding the cookie faster). This sex-specific difference in retrieval times occurred after the pre-treatment baseline (i.e. when all mice were naïve to the task), which may involve some component of learning or familiarity. The variance associated with individual subjects was comparatively high, with Subject ID accounting for 47.77% of total model variance (Estimate = 25,913.211, SE = 7,837.078, 95% CI: [10,552.82, 41,273.60], Wald *p* = 0.0009), indicating that individual mice have different performance characteristics in this metric. The binary retrieval success metric, a crude but common measurement in this assay, revealed no additional effects beyond those of the retrieval time. We observed no main effect of sex (linear mixture model *F*(1, 144) = 0.020, *p* = 0.888), time point (*F*(3, 144) = 0.014, *p* = 0.998), or treatment group (*F*(1, 144) = 0.001, *p* = 0.981). As was the case with the retrieval time, we observed a significant contribution of Subject ID (individual animal) to retrieval success (Estimate = 2.821, SE = 1.174, 95% CI: [0.520, 5.122], Wald *p* = 0.016), indicating that this metric was highly dependent on the individual animals performing the task.

**Figure 4.**
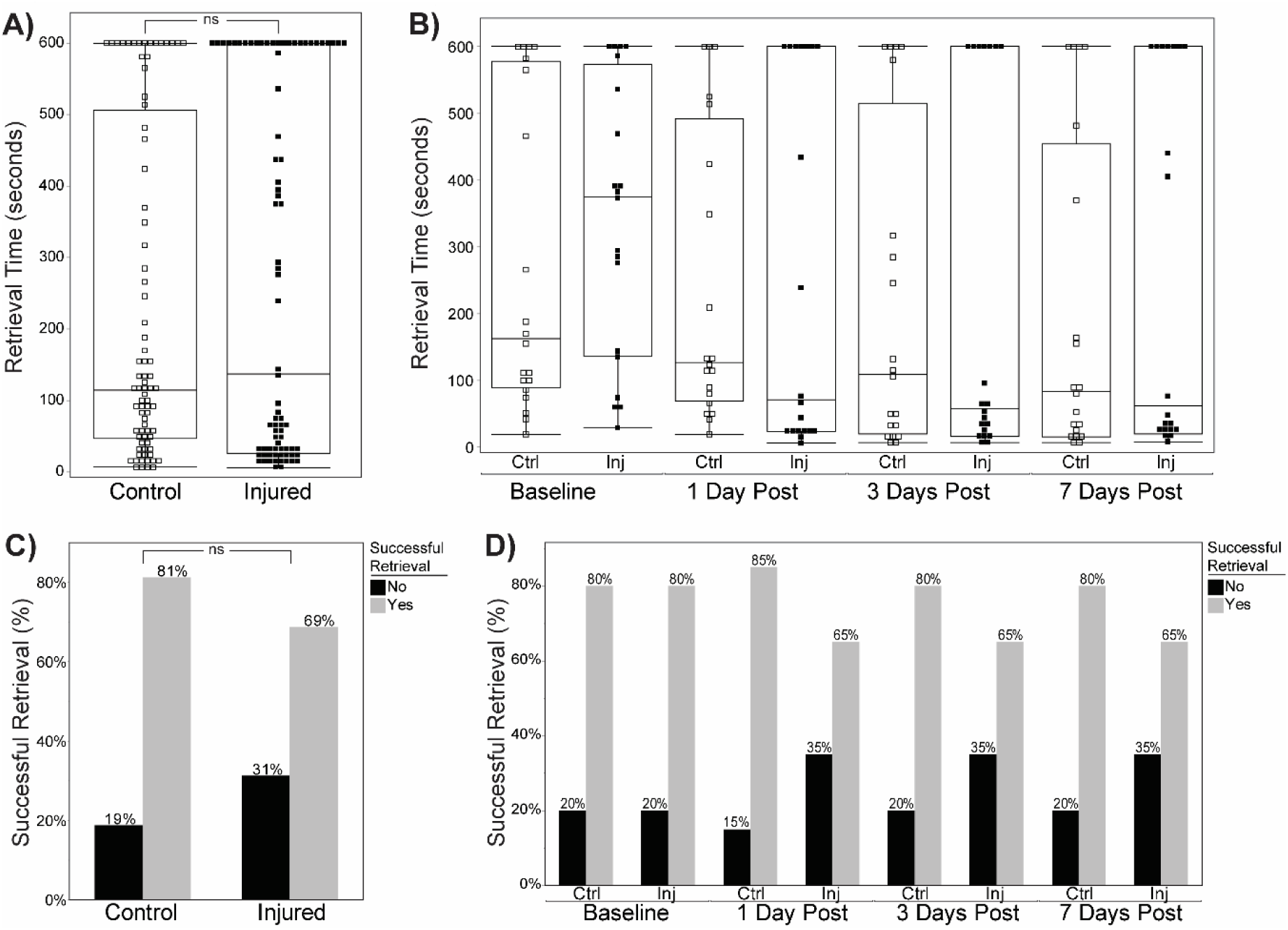
Performance on the Buried Food Task following single injury. **(A, B)** Retrieval time (in seconds) during the buried food task. **(A)** Comparison between control and injured animals averaged across all timepoints shows no significant difference in retrieval time. **(B)** Retrieval times separated by follow-up show no significant differences between control (Ctrl) and injured (Inj) groups at any timepoint. Horizontal bars indicate medians; boxes show interquartile ranges; individual points represent subjects. **(C, D)** Percentage of animals with successful food retrieval. **(C)** No significant difference in the overall proportion of successful retrievals between control and injured groups across all timepoints. **(D)** Percentage of successful retrievals by treatment group and timepoint. Control animals consistently showed higher success rates (80 – 85%) compared to injured animals (65%), no significant differences were found at any timepoint.

In the nine-impact paradigm, we evaluated retrieval time and success in the buried food tasks at regular intervals during the mTBI sequence (1 day after the 3^rd^, 6^th^, and 9^th^ impacts), and again at 1 week and 1 month following the 9^th^ impact (Fig. 5). In contrast to the results from the single-impact paradigm, we observed main effect of treatment (linear mixture model *F*(1, 4.6) = 12.613, *p* = 0.019), with increased retrieval times in the injured group compared to controls. This indicates that unlike the single-impact paradigm, rmTBI induces behavioral changes in the buried food task consistent with reduced olfactory function. Importantly, because this was a repeated test, this result could also indicate reduced attention to the task, motivation to seek a food reward, or inability to improve performance based on experience. We also observed a main effect of timepoint (linear mixture model *F*(5, 23.4) = 3.187, *p* = 0.025). Unlike the single-impact paradigm results, we did not observe a significant main effect of sex (*F*(1, 4.6) = 3.999, *p* = 0.107). We did not observe a significant interaction between sex and treatment (*F*(1, 4.6) = 1.731, *p* = 0.250), sex and timepoint (*F*(5, 23.4) = 2.130, *p* = 0.097), treatment group and timepoint (*F*(5, 23.4) = 0.938, *p* = 0.475), or an interaction between all three variables (*F*(5, 23.4) = 1.127, *p* = 0.374). Again, unlike the single-impact paradigm, the variance attributable to Subject ID was not significant, accounting for < 0.1 % of variance (Estimate = −1,062.287, SE = 3,551.960, 95% CI: [−8,676.669, 6,552.094], Wald *p* = 0.785). The simplified binary retrieval success metric provided no additional indication of significant effects of any variable (treatment, sex, timepoint) or interaction between them, demonstrating the limited utility of this measurement in the buried food test. Cumulatively, the results of this simple, limited, but accessible behavioral task were informative. They indicated that the single-impact mTBI paradigm does not cause deficits in retrieval tasks but does reveal sex-specific differences in performance when the task is repeated multiple times within a week. In the nine-impact paradigm, there were clear differences in task performance in both sexes that depended on treatment and timepoint, with rmTBI-treated animals demonstrating poorer performance (longer retrieval times) than controls.

**Figure 5.**
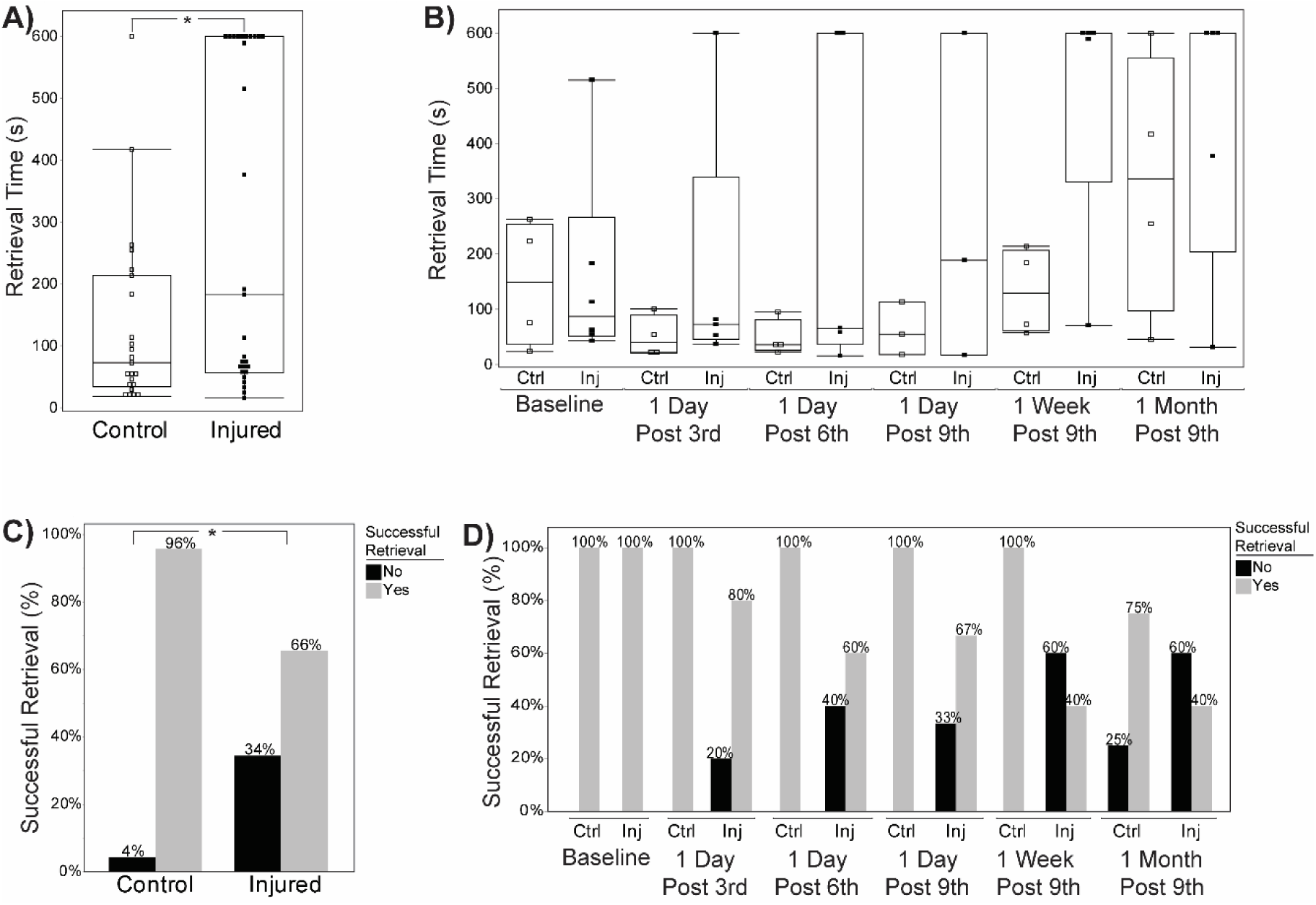
Buried Food Task performance following rmTBI in the 9-hit model. **(A, B)** Retrieval time (in seconds) during the buried food task. **(A)** Averaged across all timepoints, injured animals showed significantly longer retrieval times (*p* = 0.006). **(B)** Retrieval times separated by follow-up show no significant differences between control (Ctrl) and injured (Inj) groups at any timepoint. Horizontal bars indicate medians; boxes show interquartile ranges; individual points represent subjects. **(C, D)** Percentage of animals with successful food retrieval. **(C)** Control animals showed significantly higher rates of successful retrieval when compared to injured animals across all timepoints (*p* = 0.014). **(D)** Percentage of successful retrievals by treatment group and timepoint. Control animals consistently showed higher success rates after baseline (75 – 100%) compared to injured animals (40 – 80%), no significant differences were found at any timepoint.

### Evaluating neuroinflammation in single- and nine-impact mTBI paradigms

TBIs induce neuroinflammation in many paradigms, an indication of damage to neurons, glia, or a breakdown of the blood-brain barrier (Ziebell and Morganti-Kossmann, 2010; Logsdon et al., 2015). We therefore evaluated neuroinflammation in the brains of mice exposed to both the single-impact (Fig. 6) and nine-impact (Fig. 8) paradigms, in each case harvesting brains within 1 week of the final behavioral timepoint. We evaluated neuroinflammation via fluorescent immunohistochemical analysis of microglial markers of inflammation, Iba1 and CD68, both commonly used for this purpose (Jurga et al., 2020; Paolicelli et al., 2022; Swanson et al., 2023). We specifically quantified the abundance of low-intensity Iba1^+^ (resting microglial) somas, high-intensity Iba1^+^ (activated microglial) somas, and CD68^+^ (highly reactive, likely phagocytic) microglia (Kenkhuis et al., 2022; Wittekindt et al., 2022). To improve our ability to identify these signals across samples and brain regions and samples with differences in tissue quality we utilized pixel-based machine learning via the ilastik software platform (Berg et al., 2019). We utilized generalized linear model analysis to evaluate the abundance of these neuroinflammatory markers. The immunohistochemical analysis results for the single-impact paradigm are presented in Figures 6, 7, and Table 1 with results for the nine-impact paradigm presented in Figure 8, 9, and Table 2.

**Figure 6.**
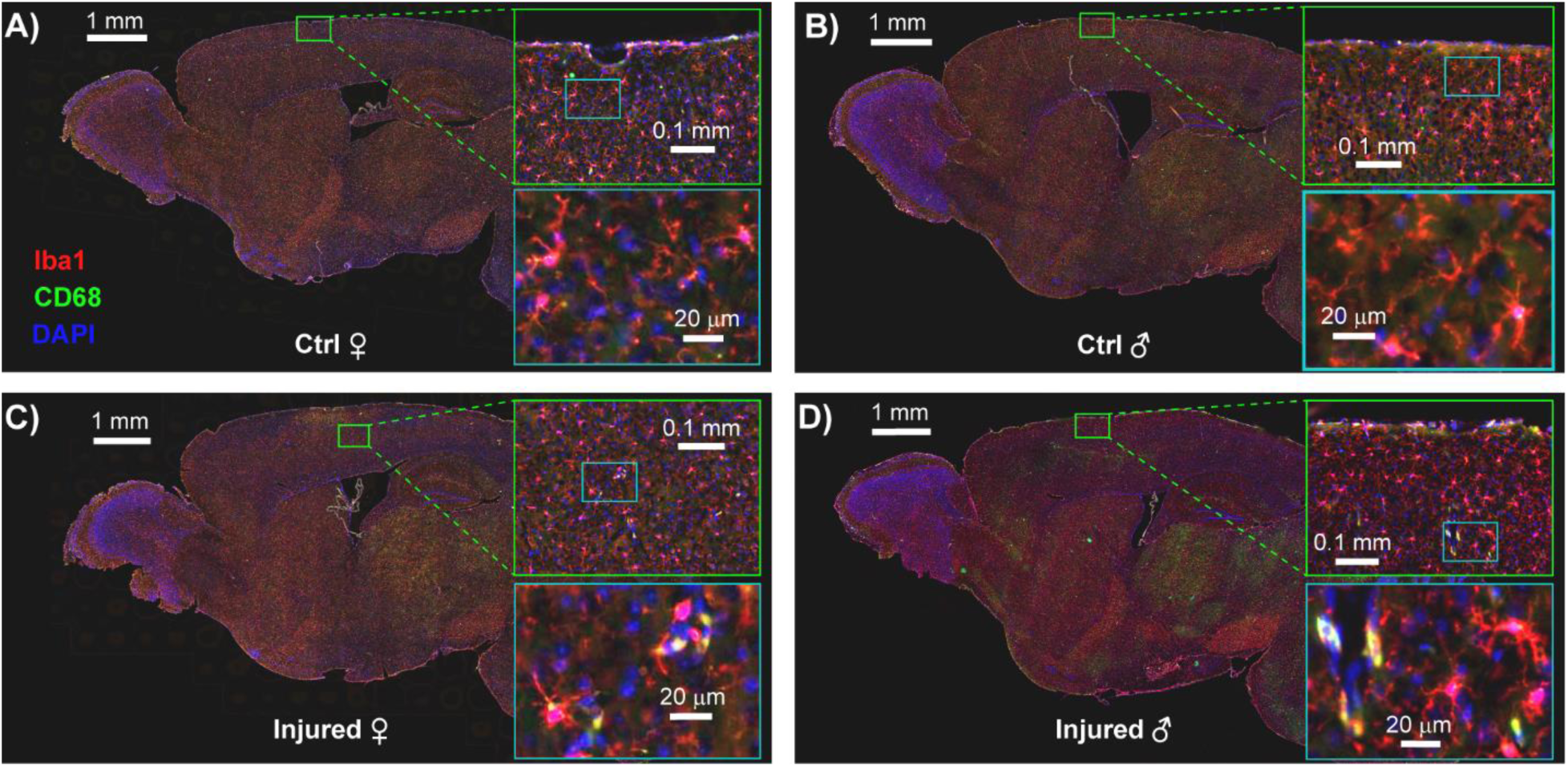
Representative IHC staining of cortical microglia across sex and injury conditions in a single mTBI model. Fluorescent immunohistochemistry (IHC) images showing Iba1⁺ (red), CD68⁺ (green), and DAPI (blue) staining in sagittal brain sections from control and injured mice, separated by sex. Panels represent a control female **(A)**, control male **(B)**, injured female **(C)**, and injured male **(D)**. Each panel includes a whole-brain sagittal section, with two sequentially magnified insets highlighting microglial staining in a defined cortical region (green box). The first inset provides intermediate magnification, while the second (light blue box) shows high magnification of the boxed region within the first. Iba1⁺ staining marks total microglia, CD68⁺ indicates lysosomal activation, and co-localization reflects reactive or phagocytic microglia. DAPI labels all nuclei. Increased Iba1⁺ and CD68⁺ signal intensity is evident in male brains—particularly injured males **(D)**—compared to females, consistent with observed sex differences in cortical microglial activation. Arrows represent approximate site of impact.

**Figure 7.**
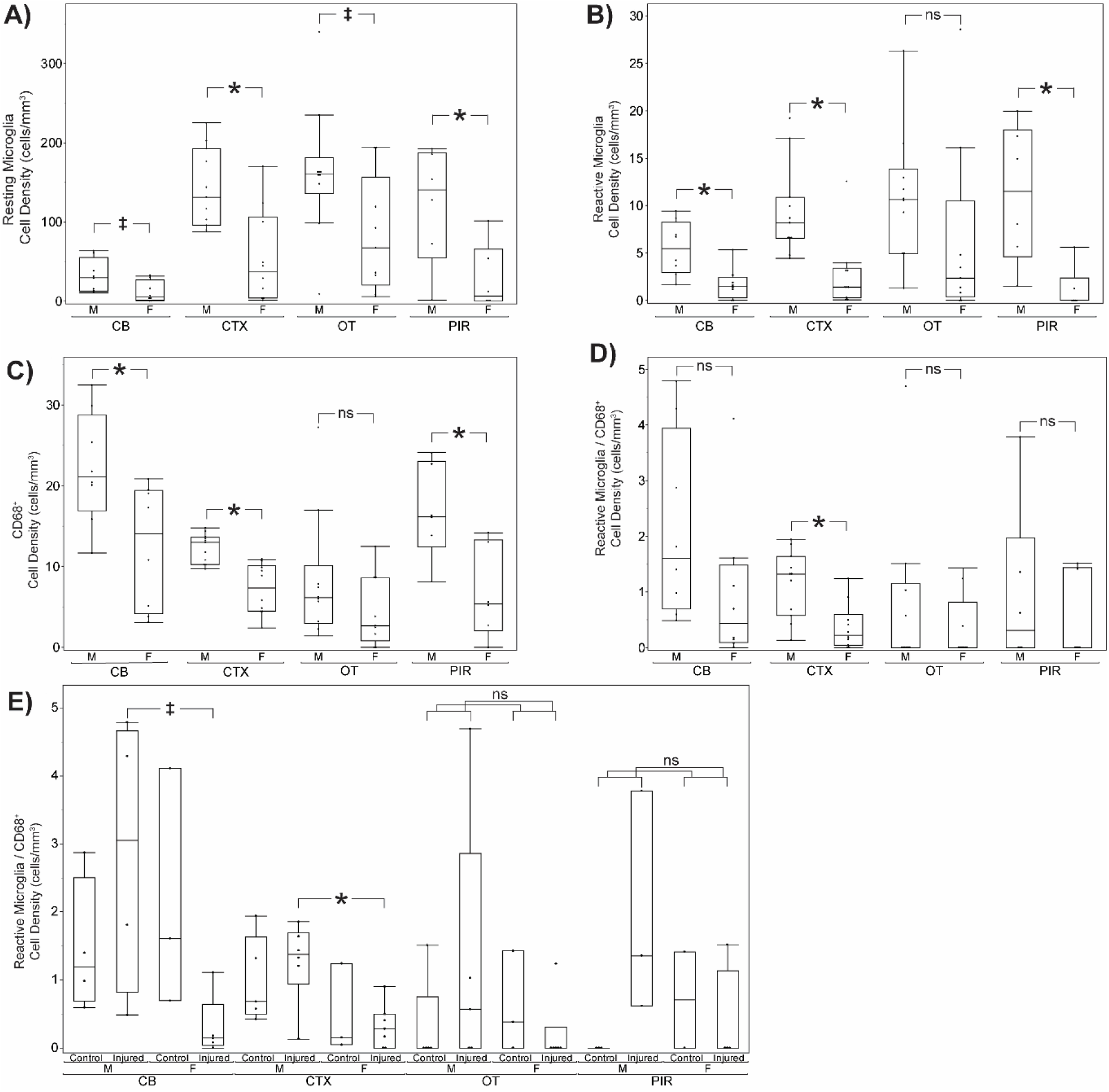
Quantitative analysis of microglial subpopulations by sex, region, and injury status. Box plots showing microglial cell density and phenotype across the cerebellum (CB), cortex (CTX), olfactory tubercle (OT), and piriform cortex (PIR), separated by sex (M = male, F = female) and treatment (Control vs. Injured). **(A)** Resting Iba1+ microglial cell density. **(B)** Reactive microglial cell density. **(C)** CD68+ (phagocytic) microglial cell density. **(D)** Dual-labeled reactive/CD68^+^ microglia. **(E)** Density of dual-labeled reactive/CD68^+^ microglia by treatment and sex. Asterisks (*) denote significant differences (*p < 0.05; ‡p < 0.10); ns = not significant.

**Figure 8.**
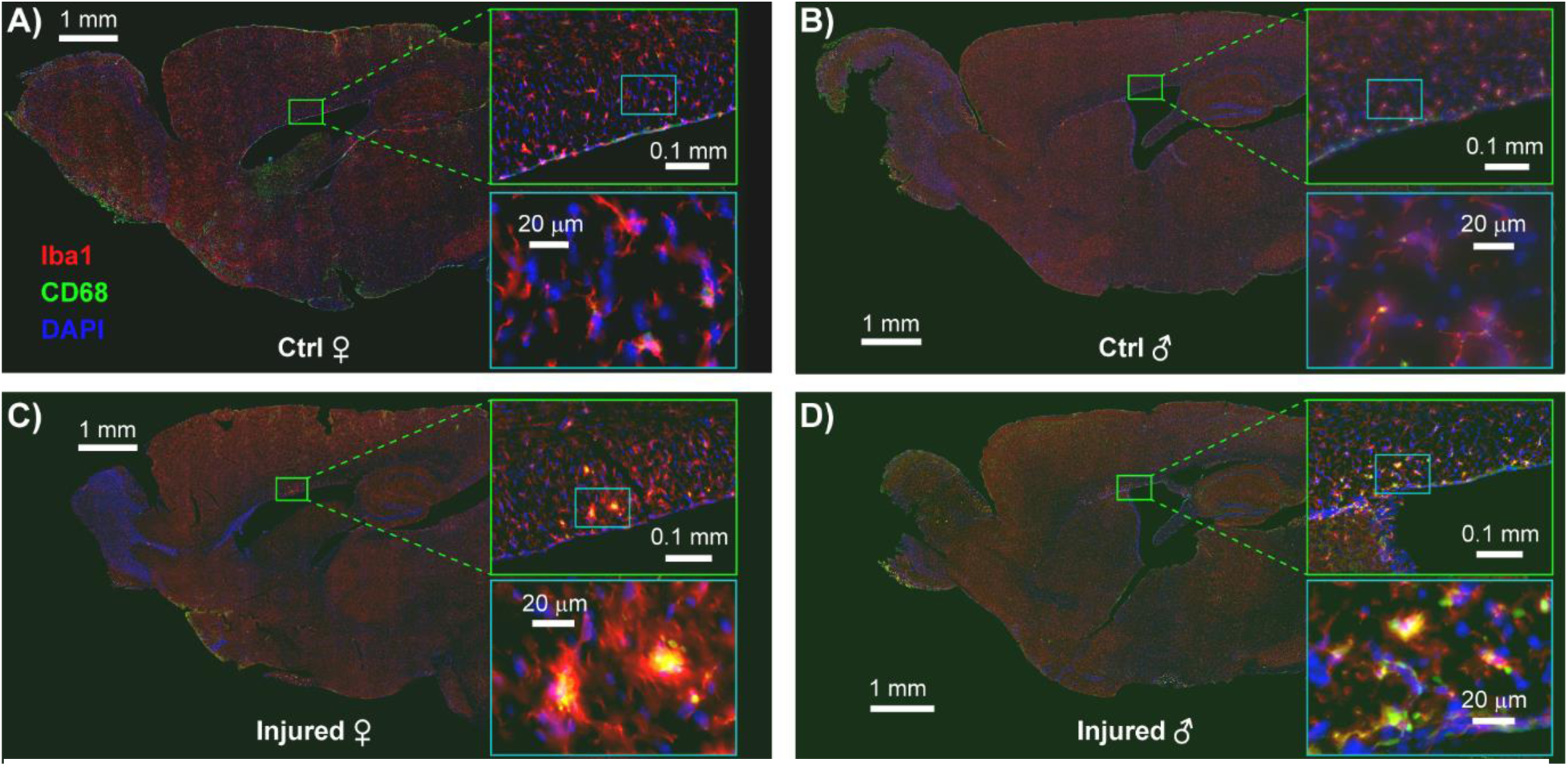
Representative IHC staining of corpus callosum microglial activation across experimental groups in rmTBI model. Sagittal brain slices were stained for Iba1 (red; microglia), CD68 (green; activated/phagocytic microglia), and DAPI (blue; nuclei). Each panel shows a full sagittal section with the corpus callosum highlighted in zoomed-in insets: the upper inset shows a regional overview, and the lower inset shows high-magnification views of individual microglial profiles. **(A)** Control female, **(B)** Control male, **(C)** Injured female, **(D)** Injured male. Increased CD68 signal intensity and altered microglial morphology are evident in the injured female and male brain **(C, D)**, indicating enhanced microglial activation following rmTBI.

**Figure 9.**
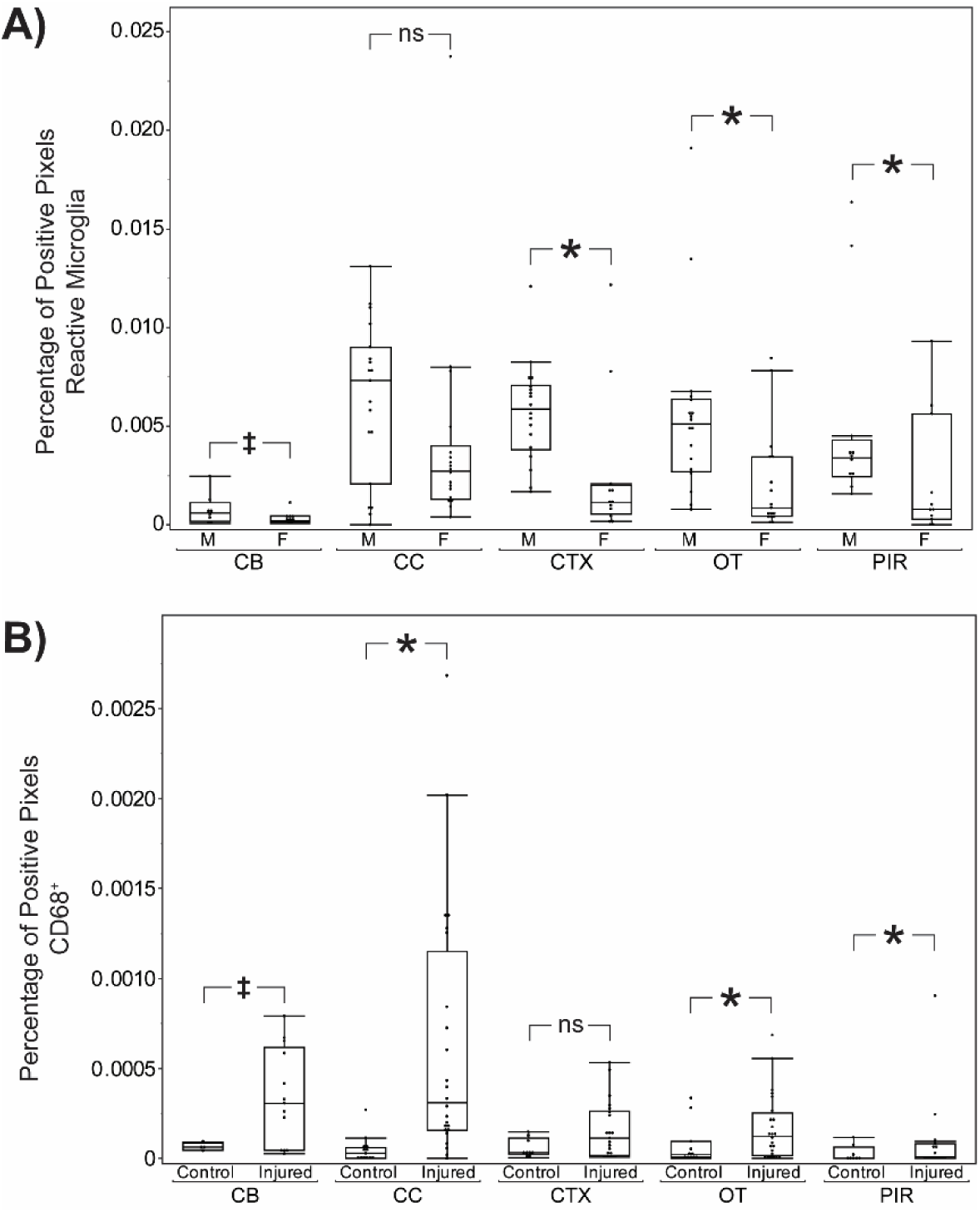
Quantification of reactive and phagocytic microglia across brain regions and treatment groups. **(A)** Percentage of positive pixels for reactive microglia across brain regions, separated by sex and treatment condition. **(B)** Percentage of CD68-positive pixels, representing phagocytic microglia, across the same groups and regions. Significant differences are indicated by asterisks (*p < 0.05); trend-level differences (†) and non-significant comparisons (ns) are also noted where appropriate.

**Table 1.** Two-way ANOVA Results for Single Impact TBI Model.

| Term/Marker | AOB | CB | CTX | HB | HPF | MOB | OT | PIR | SN |
| --- | --- | --- | --- | --- | --- | --- | --- | --- | --- |
| Sex/Resting Microglia | **0.031** | 0.055 | **0.006** | **0.001** | **0.014** | **0.005** | 0.057 | **0.019** | **0.003** |
|  | ♀ ↓ | ♀ ↓ | ♀ ↓ | ♀ ↓ | ♀ ↓ | ♀ ↓ | ♀ ↓ | ♀ ↓ | ♀ ↓ |
| Treatment/Resting Microglia | 0.664 | 0.539 | 0.561 | 0.143 | 0.948 | 0.772 | 0.781 | 0.214 | 0.948 |
| Sex*Treatment/Resting Microglia | 0.136 | 0.747 | 0.158 | 0.257 | 0.503 | 0.074 | **0.018** | 0.154 | 0.335 |
|  |  |  |  |  |  | †♂ ↑ | †♂ ↑ |  |  |
| Sex/Reactive Microglia | 0.091 | **0.004** | **0.005** | **0.017** | 0.136 | 0.076 | 0.414 | **0.015** | **0.009** |
|  | ♀ ↓ | ♀ ↓ | ♀ ↓ | ♀ ↓ |  | ♀ ↓ |  | ♀ ↓ | ♀ ↓ |
| Treatment/Reactive Microglia | 0.888 | 0.170 | 0.901 | 0.166 | 0.552 | 0.286 | 0.759 | 0.374 | 0.877 |
| Sex*Treatment/Reactive Microglia | 0.302 | 0.086 | 0.805 | 0.419 | 0.516 | 0.206 | 0.121 | 0.681 | 0.949 |
|  |  | †♂ ↓ |  |  |  |  |  |  |  |
| Sex/CD68+ | 0.848 | **0.033** | **0.000** | **0.006** | **0.020** | 0.321 | 0.221 | **0.034** | **0.034** |
|  |  | ♀ ↓ | ♀ ↓ | ♀ ↓ | ♀ ↓ |  |  | ♀ ↓ | ♀ ↓ |
| Treatment/CD68+ | 0.483 | 0.938 | 0.467 | 0.442 | 0.495 | 0.189 | 0.454 | 0.364 | 0.658 |
| Sex*Treatment/CD68+ | 0.487 | 0.769 | 0.969 | 0.354 | 0.911 | 0.669 | 0.555 | 0.753 | 0.466 |
| Sex/Double Positive | 0.108 | 0.206 | **0.009** | 0.075 | **0.008** | 0.053 | 0.615 | 0.508 | 0.062 |
|  |  |  | ♀ ↓ | ♀ ↓ | ♀ ↓ | ♀ ↓ |  |  | ♀ ↓ |
| TBI/Double Positive | 0.847 | 0.749 | 0.811 | 0.931 | 0.648 | 0.658 | 0.501 | 0.222 | 0.475 |
| Sex*Treatment/Double Positive | 0.924 | **0.040** | 0.390 | 0.153 | 0.424 | 0.914 | 0.232 | 0.099 | 0.498 |
|  |  | †♂ ↑ |  |  |  |  |  | †♂ ↑ |  |

*p*-values from two-way ANOVA testing of the effects of sex, treatment, and their interaction on Iba1<sup>+</sup>, CD68<sup>+</sup>, and Iba1<sup>+</sup>/CD68<sup>+</sup> co-labeled in different ROIs. Each row represents an ANOVA term (main or interaction effect) for a specific microglia marker or cell type (resting, reactive, CD68<sup>+</sup>, double positive). Values in **red** indicate statistically significant effects ( $p < 0.05$ ). † = Injured, ♂ = Male, and ♀ = Female. Direction of arrow represents direction of effect.

**Table 2.** Results from Generalized Linear Mixed Model (GLMM) analyses in the Nine-Impact rmTBI Model.

| Term/Marker | ACB | AOB | AON | CB | CC | CP | CTX | HB | HPF | LOT | MOB | OT | PIR | SN |
| --- | --- | --- | --- | --- | --- | --- | --- | --- | --- | --- | --- | --- | --- | --- |
| Sex/Resting Microglia | 0.369 | 0.661 | 0.458 | 0.068 | 0.517 | 0.419 | 0.415 | 0.330 | 0.301 | 0.423 | 0.627 | 0.353 | 0.385 | 0.775 |
|  |  |  |  | ♀ ↓ |  |  |  |  |  |  |  |  |  |  |
| Treatment/Resting Microglia | 0.642 | 0.395 | 0.353 | 0.652 | 0.750 | 0.708 | 0.783 | 0.661 | 0.474 | 0.644 | 0.640 | 0.941 | 0.903 | 0.833 |
| Sex*Treatment/Resting Microglia | 0.438 | 0.628 | 0.281 | 0.668 | 0.353 | 0.501 | 0.468 | 0.465 | 0.360 | 0.136 | 0.978 | 0.627 | 0.662 | 0.296 |
| Sex/Reactive Microglia | 0.044 | 0.037 | 0.048 | 0.071 | 0.228 | 0.081 | 0.006 | 0.031 | 0.003 | 0.061 | 0.020 | 0.031 | 0.036 | 0.142 |
|  | ♀ ↓ | ♀ ↓ | ♀ ↓ | ♀ ↓ |  | ♀ ↓ | ♀ ↓ | ♀ ↓ | ♀ ↓ | ♀ ↓ | ♀ ↓ | ♀ ↓ | ♀ ↓ |  |
| Treatment/Reactive Microglia | 0.680 | 0.524 | 0.734 | 0.361 | 0.482 | 0.837 | 0.102 | 0.363 | 0.593 | 0.829 | 0.806 | 0.219 | 0.319 | 0.716 |
| Sex*Treatment/Reactive Microglia | 0.275 | 0.254 | 0.395 | 0.229 | 0.959 | 0.499 | 0.062 | 0.178 | 0.107 | 0.688 | 0.681 | 0.128 | 0.087 | 0.444 |
|  |  |  |  |  |  |  | †♀ ↑ |  |  |  |  |  | †♂ ↓ |  |
| Sex/Microglial Processes | 0.424 | 0.411 | 0.280 | 0.422 | 0.613 | 0.472 | 0.272 | 0.430 | 0.234 | 0.355 | 0.246 | 0.294 | 0.357 | 0.640 |
| Treatment/Microglial Processes | 0.719 | 0.523 | 0.801 | 0.786 | 0.696 | 0.697 | 0.900 | 0.764 | 0.341 | 0.701 | 0.826 | 0.702 | 0.680 | 0.743 |
| Sex*Treatment/Microglial Processes | 0.628 | 0.353 | 0.449 | 0.589 | 0.319 | 0.530 | 0.644 | 0.519 | 0.295 | 0.469 | 0.292 | 0.905 | 0.951 | 0.418 |
| Sex/CD68+ | 0.962 | 0.126 | 0.481 | 0.566 | 0.621 | 0.957 | 0.757 | 0.690 | 0.274 | 0.981 | 0.651 | 0.190 | <.001 | 0.512 |
|  |  |  |  |  |  |  |  |  |  |  |  |  | ♀ ↓ |  |
| Treatment/CD68+ | 0.254 | 0.105 | 0.350 | 0.064 | 0.012 | 0.034 | 0.102 | 0.049 | 0.093 | 0.386 | 0.607 | 0.049 | <.001 | 0.458 |
|  |  |  |  | † ↑ | † ↑ | † ↑ |  | † ↑ | † ↑ |  |  | † ↑ | † ↑ |  |
| Sex*Treatment/CD68+ | 0.245 | 0.775 | 0.400 | 0.407 | 0.306 | 0.174 | 0.186 | 0.281 | 0.079 | 0.878 | 0.390 | 0.287 | <.001 | 0.817 |
|  |  |  |  |  |  |  |  |  | †♀ ↑ |  |  |  | †♂ ↑ |  |

*p*-values from GLMM analyses testing the effects of sex, treatment, and their interaction on low-intensity (resting) Iba1<sup>+</sup> somas, high-intensity (reactive) Iba1<sup>+</sup> somas, Iba1<sup>+</sup> microglial processes, and CD68<sup>+</sup> puncta across different ROIs. Each row represents an ANOVA term (main or interaction effect) for a specific microglial marker or cell type (resting, reactive, processes, or CD68<sup>+</sup>). Values in red indicate statistically significant effects ( $p < 0.05$ ). *Note:* A general *p*-value was included in CD68<sup>+</sup> results from the PIR because the model identified complete separation, preventing an exact estimate of variance and significance. † = Injured, ♂ = Male, and ♀ = Female. Direction of arrow represents direction of effect. **Abbreviations:** ACB = nucleus accumbens, AOB = accessory olfactory bulb, AON = anterior olfactory nucleus, CB = cerebellum, CC = corpus callosum, CP = caudoputamen, CTX = cortex, HB = hindbrain, HPF = hippocampal formation, LOT = lateral olfactory tract, MOB = main olfactory bulb, OT = olfactory tubercle, PIR = piriform cortex, SN = substantia nigra

### Sex-specific differences in microglial activation following single mTBI

In the cortex, male mice consistently exhibited significantly greater microglial presence and activation than females across all markers assessed. Significant sex differences were observed for Iba1⁺ resting microglia staining (141.08 ± 16.13 in males vs. 64.25 ± 18.39 in females; two-way ANOVA, *F*(1, 17) = 9.865, *p* = 0.006), Iba1⁺ reactive microglia (9.40 ± 1.37 vs. 2.71 ± 1.57; two-way ANOVA *F*(1, 17) = 10.319, *p* = 0.005), CD68⁺ cells (12.24 ± 0.79 vs. 7.01 ± 0.90; two-way ANOVA *F*(1, 17) = 19.113, *p* = 0.0004), and Iba1⁺/CD68⁺ double-positive microglia (1.13 ± 0.16 vs. 0.40 ± 0.19; two-way ANOVA *F*(1, 17) = 8.693, *p* = 0.009). These results indicate a robust and consistent sex-specific increase in cortical microglial activation in male mice, independent of injury.

Outside of the cortex, similar patterns emerged, though varied across regions and cell types. In the AOB, males (N = 10) again showed significantly higher levels of Iba1^+^ resting microglia staining versus females (N = 8) (138.14 ± 22.13 vs. 56.93 ± 25.55; two-way ANOVA *F*(1, 14) = 5.772, *p* = 0.031).

Analysis of the CB revealed males (N = 8) exhibited significantly greater levels of Iba1^+^ reactive microglia staining versus females (N = 8) (5.50 ± 0.73 vs. 1.70 ± 0.76; two-way ANOVA *F*(1, 12) = 13.056, *p* = 0.004). Additionally, an interaction between sex and treatment was observed for double-positive cells in the CB, with injured males (N = 4) exhibiting the highest levels of co-labeled microglia and injured females showing a marked reduction (2.84 ± 0.69 vs. 0.30 ± 0.61; two-way ANOVA *F*(1, 12) = 5.329, *p* = 0.040).

In the HB, we observed males (N = 9) exhibiting higher staining levels than females (N = 8) in Iba1^+^ resting microglia (96.61 ± 10.46 vs. 34.42 ± 11.38; two-way ANOVA *F*(1, 13) = 16.192, *p* = 0.001), Iba1^+^ reactive microglia (8.26 ± 1.45 vs. 2.39 ± 1.58; two-way ANOVA *F*(1, 13) = 7.485, *p* = 0.017), and CD68^+^ cells (14.58 ± 1.10 vs. 9.20 ± 1.20; two-way ANOVA *F*(1, 13) = 10.970, *p* = 0.006).

Similarly, in the HPF we observed elevated levels of staining for males (N = 9) when compared to females (N = 9) in Iba1^+^ resting microglia (109.17 ± 17.06 vs. 39.71 ± 17.99; two-way ANOVA *F*(1, 14) = 7.850, *p* = 0.040), CD68^+^ cells (16.25 ± 2.00 vs. 8.65 ± 2.10; two-way ANOVA *F*(1, 14) = 6.858, *p* = 0.020), and Iba1^+^/CD68^+^ double-positive cells (1.85 ± 0.30 vs. 0.48 ± 0.32; two-way ANOVA *F*(1, 14) = 9.763, *p* = 0.008). In the MOB, males (N = 10) again displayed significantly higher levels of Iba1^+^ resting microglia when compared to female mice (N = 8) (135.71 ± 12.73 vs. 70.43 ± 14.70; two-way ANOVA *F*(1, 14) = 11.264, *p* = 0.005).

When analyzing staining in the OT, we found a significant interaction between sex and treatment for Iba^+^ resting microglia (two-way ANOVA *F*(1, 15) = 7.107, *p* = 0.018). Intriguingly, Tukey HSD post hoc comparisons revealed a sex-dependent bidirectional shift following injury with injured males (N = 5) exhibiting an increase compared to control males (N = 5) (212.13 ± 30.88 vs. 115.51 ± 30.88) and injured females (N = 6) exhibiting a decrease compared to control females (N = 3) (57.22 ± 28.19 vs. 135.25 ± 28.19).

In the PIR, we again observed male mice (N = 6) demonstrating significantly higher levels of staining when compared to females in Iba1^+^ resting microglia (122.06 ± 21.69 vs. 29.21 ± 23.01; two-way ANOVA *F*(1, 8) = 8.623, *p* = 0.019), Iba1^+^ reactive microglia (11.24 ± 2.29 vs. 0.86 ± 2.43; two-way ANOVA *F*(1, 8) = 9.632, *p* = 0.015), and CD68^+^ cells (16.85 ± 2.48 vs. 7.58 ± 2.63; two-way ANOVA *F*(1, 8) = 6.569, *p* = 0.034). This may suggest widespread sex-specific activation in this olfactory processing region.

Finally, in the SN, we again observed males (N = 9) showing greater staining when compared to females (N = 8) across Iba1^+^ resting microglia (292.69 ± 35.90 vs. 101.48 ± 39.08; two-way ANOVA *F*(1, 13) = 12.983, *p* = 0.003), Iba1^+^ reactive microglia (32.55 ± 6.51 vs. 3.28 ± 7.09; two-way ANOVA *F*(1, 13) = 9.253, *p* = 0.009), and CD68^+^ cells (17.70 ± 2.86 vs. 7.71 ± 3.11; two-way ANOVA *F*(1, 13) = 5.603, *p* = 0.034).

### Sex- and treatment-specific differences in microglial activation and lysosomal activity in a nine-impact rmTBI model

To complement the findings of the single-impact model, we conducted an independent analysis utilizing ilastik to binarize IHC images from the nine-impact rmTBI model using a generalized linear mixed model (GLMM) with a binomial distribution and logit link. This approach evaluated the proportion of high-intensity pixels representing positive staining across brain regions to determine if sex, treatment, or the interaction between sex and treatment had a significant effect on the presence of positively stained pixels. Additionally, subject ID was included as a random effect to control for multiple tissue sections coming from the same animal.

Sex effects were prominent across multiple olfactory and limbic regions, with male mice generally exhibiting greater microglial activation than females. In the ACB, males (N = 17) showed significantly higher odds of high-intensity Iba1⁺ staining compared to females (N = 15), with an estimated odds ratio of 10.00 ± 8.59 (95% CI: [1.10, 90.96]; *F*(1, 5) = 7.192, *p* = 0.044). Similar effects were observed in the AOB with males (N = 5) again showing significantly higher odds of high-intensity Iba1^+^ staining when compared to females (N = 5) with an estimated odds ratio of 6.75 ± 3.61 (95% CI: [1.23, 36.94]; *F*(1, 3) = 12.775, *p* = 0.037) and in the AON with males (N = 16) having an estimate odds ratio of 7.65 ± 5.99 (95% CI: [1.02, 57.23]; *F*(1, 5) = 6.748, *p* = 0.048) when compared to females (N = 16). Similar reductions in high-intensity Iba1^+^ microglial somas were seen in cortical regions with male mice again exhibiting significantly higher odds in the CTX (N = 18 males and 12 females, Odds Ratio = 12.22 ± 6.68, 95% CI: [3.00, 49.78]; *F*(1, 5) = 20.967, *p* = 0.006), HPF (N = 15 males and 13 females, Odds Ratio = 10.54 ± 4.53, 95% CI: [3.49, 31.80]; *F*(1, 5) = 30.054, *p* = 0.003), and MOB (N = 13 males and 5 females, Odds Ratio = 6.72 ± 2.84, 95% CI: [1.76, 25.73]; *F*(1, 3) = 20.423, *p* = 0.020). We also observed further sex-dependent differences in microglial activation with males displaying significantly higher odds of high-intensity Iba1^+^ staining in the HB (N = 11 males and 11 females, Odds Ratio = 5.48 ± 2.86, 95% CI: [1.29, 23.33]; *F*(1, 4) = 10.630, *p* = 0.031), OT (N = 18 males and 17 females, Odds Ratio = 8.11 ± 5.70, 95% CI: [1.33, 49.33]; *F*(1, 5) = 8.881, *p* = 0.031), and PIR (N = 12 males and 11 females, Odds Ratio = 7.74 ± 5.56, 95% CI: [1.22, 49.09]; *F*(1, 5) = 8.098, *p* = 0.036). These binarized GLMM-based findings from the nine-impact model confirm and extend previous findings from traditional cell count analysis in the single-impact cohort, supporting a robust sex effect in microglial activation patterns.

Treatment effects were most apparent in CD68^+^ puncta staining (Fig. 9B), indicating injury-related microglial lysosomal activation. In the CC, we observed injured mice (N = 24) exhibited significantly greater odds of positively identified staining when compared to control mice (N = 13) (Odds Ratio = 12.71 ± 8.34, 95% CI: [2.36, 68.47]; *F*(1, 5) = 15.028, *p* = 0.012). We did not observe a significant effect of sex (*p* = 0.621) or a significant interaction between sex and treatment (*p* = 0.306) suggesting this increase in CD68^+^ is primarily driven by injury status rather than sex. We observed similar treatment effects in the CP (N = 19 injured and 8 control, Odds Ratio = 15.99 ± 15.30, 95% CI: [1.38, 185.82]; *F*(1, 5.1) = 8.382, *p* = 0.034), HB (N = 16 injured and 6 control, Odds Ratio = 10.90 ± 9.33, 95% CI: [1.01, 117.33]; *F*(1, 4) = 7.779, *p* = 0.049), and OT (N = 24 injured and 11 control, Odds Ratio = 7.11 ± 5.39, 95% CI: [1.01, 49.83]; *F*(1, 5) = 6.678, *p* = 0.049). These results suggest that rmTBI leads to increased lysosomal activity across multiple subcortical regions involved in sensorimotor and limbic processing.

## Discussion

In this study, we developed and validated a novel weight-drop apparatus capable of inducing closed-skull repetitive mild traumatic brain injury (rmTBI) in mice while capturing high-fidelity biomechanical data. This custom-engineered system allowed for precise delivery of impacts with controlled velocity and force, real-time accelerometry, and high-speed video capture of head kinematics. Coupled with behavioral and histological analyses, this model provides a robust platform to investigate the mechanistic underpinnings and long-term consequences of rmTBI.

### Creating an instrument with reliable impact forces and quantifiable kinematic effects

Across both single and repetitive impact paradigms, we observed low inter-subject variability in impact velocity and force, demonstrating that the apparatus delivers reproducible mechanical performance. The negligible variance attributed to Subject ID in our statistical tests underscores the consistency of impact delivery. Given that reproducibility can be a source of concern in closed skull mTBI models, this is a major strength of our instrument and approach. Any laboratory adopting this approach should note that the performance of the instrument depends on the specific materials used (including fasteners, etc.). It is critical that each instrument (and updates to it) be evaluated, and measurements of impact forces produced, to ensure that the impacts delivered remain in the proper range for mTBI production.

We strongly recommend that any laboratory adopting this method install the rebound-arresting solenoid system, which introduces stronger whiplash components that contribute strongly to TBI sequelae (Jang and Kwon, 2018; Al-Khazali et al., 2022; Belhassen et al., 2023). This system additionally eliminates unintended secondary impacts, allowing isolation of the primary biomechanical insult. We also strongly recommend the implementation of high-speed video monitoring of every impact. This is important because even with consistent instrument performance, the response of each animal to the impact varies based on the exact positioning of the head under the impactor, the animal’s sex, its age, and other factors. By recording a high-speed video of each impact, one can quantify of both linear and angular accelerations to ensure that the experiences of each animal are consistent. This also allows users to catch cases in which higher or lower acceleration was experienced by an animal than the average. In theory, these measurements could be useful when evaluating correlations between kinematic factors and other behavioral and structural measurements. In this initial set of experiments, for example, we observed statistically significant variability in upward linear acceleration across impact sessions, particularly at Impacts #4, #7, and #9. Interestingly, both upward and downward angular accelerations peaked at Impact #2, suggesting a transient period of increased biomechanical transmission to the brain, possibly due to cumulative priming or changes in tissue compliance. Furthermore, sex differences emerged in kinematic responses, particularly in upward angular acceleration, where female mice showed reduced values. This aligns with prior studies suggesting sex-based differences in neck musculature and head-neck biomechanics may influence TBI susceptibility or symptomology (Tierney et al., 2005).

### Behavioral outcomes and regional neuroinflammation

Fully evaluating the behavioral effects of this rmTBI model will require considerable effort. We chose to perform the buried cookie task because it has components similar to the open field task (in which mice naturally explore a new arena) and includes olfaction (a focus of the laboratory) and foraging. In the single impact model, we did not observe extensive changes to retrieval time or retrieval success, suggesting that the behavioral response to a single mTBI is mild. This is perhaps unsurprising, as isolated mTBIs in humans are thought to be dramatically underreported, presumably because of the lack of obvious sustained behavioral changes in the affected individuals (Buck, 2011; Yue et al., 2023). In contrast, we measured prolonged latency to retrieve the buried food in the 9-impact model, particularly at later timepoints (Fig. 5B). This is a simple measurement and a simple behavioral assay, but the results support the conclusion that the 9-hit rmTBI model causes dysfunction in sensory, motor, and/or memory pathways, mirroring symptoms experienced by humans suffering from rmTBIs (List et al., 2015; Bielanin et al., 2024). Notably, most mice still successfully retrieved the buried food eventually, either suggesting that the rmTBI model does not induce total anosmia, or that rmTBI-affected mice can use slower, less-efficient strategies to locate buried food in the absence of the sense of smell.

Neuroinflammation is a typical consequence of mTBIs (Patterson and Holahan, 2012; Malik et al., 2023). Our immunohistochemical analyses of microglial abundance and activation state in the single and multiple impact models revealed consistent sex-based differences in the numbers of activated microglia across brain regions, with male mice exhibiting greater levels of Iba1^+^ and CD68^+^ staining. This is consistent with multiple reports of sex differences in microglial abundance and reactivity (Guneykaya et al., 2018; Bordt et al., 2020; Han et al., 2021). In the nine-impact model, we observed region-specific increases in CD68^+^ puncta in injured mice, especially in white matter-rich regions such as the corpus callosum and caudoputamen, consistent with microglial recruitment and phagocytic activation. Notably, the piriform cortex exhibited a statistical interaction between sex and treatment in both models, indicating that this important olfactory region is affected in a sex-specific manner. At this time, we interpret this result cautiously, and expect future inquiries to be needed to understand the piriform cortex’s potential vulnerability following rmTBI (Witkowski et al., 2019).

### Model Limitations and Considerations

While the modified weight-drop model demonstrates high reproducibility and offers valuable mechanistic insights, several limitations warrant consideration. First, despite the closed-skull approach and control over impact kinematics, this model does not account for individual variability in skull thickness or brain anatomy that could subtly influence impact biomechanics. Second, the use of external positioning via anatomical landmarks introduces some degree of variability in precise impact location, though the use of crosshair lasers minimized this effect.

Additionally, behavioral assays were limited to olfactory-based tasks. While the buried cookie test is a popular test for olfactory dysfunction and includes components of sensory and motor function, it does not capture other cognitive or affective impairments commonly associated with rmTBI, such as working memory deficits, anxiety-like behavior, or motor coordination challenges (Peng et al., 2016). Future studies could incorporate a broader behavioral battery to comprehensively evaluate the neurofunctional consequences of injury.

The immunohistochemical evaluation of neuroinflammation also has inherent constraints. Although ilastik-based segmentation provided objective quantification of cell-specific markers, Iba1-associated fluorescence intensity may not perfectly correlate with functional state or phagocytic activity (Green and Rowe, 2024). Moreover, the relatively small number of animals per group, particularly when stratified by sex, may limit statistical power in detecting interaction effects. These limitations underscore the need for larger-scale validation studies and the integration of complementary readouts such as transcriptomics or cytokine profiling.

### Translational Relevance

The capacity to control for both linear and angular acceleration while preventing rebound injuries addresses critical gaps in current preclinical TBI models, many of which fail to replicate the rotational forces thought to drive diffuse axonal injury and chronic neuroinflammation (Risling et al., 2019; Mayer et al., 2021; Umfress et al., 2023). By aligning our model with human-relevant mechanisms, particularly repetitive sub-concussive exposure and its cumulative effects, this platform offers a translationally relevant framework for investigating injury thresholds, temporal windows of vulnerability, and sex-specific responses.

Moreover, the sex differences observed in both kinematic responses and neuroinflammatory profiles mirror emerging human literature showing differential outcomes in males and females following mTBI (Gupte et al., 2019; Starkey et al., 2022; Baskin et al., 2023). These differences have often been underexplored or conflated in preclinical studies, and our findings provide further evidence that sex is not only a biological variable but a potential modifier of injury response.

### Future Directions

Building on this foundation, future work should focus on mapping the temporal evolution of microglial activation and its relationship to behavioral recovery or decline. Longitudinal *in vivo* imaging or CSF biomarker analyses could further elucidate the dynamic interplay between inflammation and function. Additionally, expanding behavioral phenotyping beyond olfaction to include measures of cognitive flexibility, sensorimotor gating, and social interaction will allow for a more holistic assessment of injury outcomes.

Another promising avenue lies in integrating electrophysiological recordings to assess circuit-level dysfunction following rmTBI (Witkowski et al., 2019). Given the apparent region-specific vulnerability observed in our histological data, particularly in olfactory and limbic structures, targeted recordings could reveal disruptions in synaptic plasticity or oscillatory activity that underlie behavioral impairments.

Finally, pharmacological intervention studies using this model could help identify neuroprotective agents or anti-inflammatory strategies that are effective across sexes or during specific phases of the injury cascade. By leveraging the precision and reliability of our impact apparatus, we can systematically test therapeutic timing and dose-response relationships with high translational fidelity.

### Summary

In summary, we introduce a refined model of repetitive mild traumatic brain injury that combines biomechanical precision, reproducibility, and translational relevance. Our results highlight both consistent injury delivery and biologically meaningful outcomes across behavioral and histological domains. The sex differences observed in the kinematic response to impacts, performance in the buried cookie task, and neuroinflammatory markers underscore the importance of sex-specific analyses in TBI research. This model represents a powerful tool for dissecting the mechanistic underpinnings of rmTBI and lays the groundwork for future investigations into injury progression, resilience, and therapeutic intervention.

## Acknowledgments

We thank Michael Mastrangelo for technical assistance and support of the animal colony. Images for the rmTBI IHC were taken in the URMC Center for Advanced Light Microscopy and Nanoscopy (CALMN). Partial support for the project was provided by grants from the National Institutes of Health (Grants R01DC021213 and R01DC015784), Institutional Funds from the University of Rochester School of Medicine and Dentistry, and a pilot grant from the Texas Institute for Brain Injury and Repair at University of Texas Southwestern Medical Center (all to JPM).

